# ContextTAD: Context-aware boundary learning for TAD calling from Hi-C contact maps

**DOI:** 10.64898/2026.05.08.723772

**Authors:** Weicai Long, Yusen Hou, Yanlin Zhang

## Abstract

**Motivation:** Reliable topologically associating domain (TAD) calling from Hi-C contact maps remains difficult at high resolution and realistic sequencing depth. A central reason is that many callers learn boundary evidence largely from local signals, while domain compatibility is handled mainly during downstream decoding, so the learned boundary scores are not explicitly optimized for the TAD assembly step that ultimately determines the final calls.

**Results:** We present ContextTAD, a deep-learning TAD caller that learns boundary evidence from broader local Hi-C windows that capture TAD-scale structural context. Instead of treating boundary prediction as an isolated per-bin classification problem, ContextTAD uses a context-aware representation to produce left- and right-boundary tracks that are explicitly optimized for downstream TAD assembly. Concretely, the model combines multiscale feature extraction from 2D Hi-C windows with a pair objective that rewards compatible boundary combinations and a count objective that regularizes window-level boundary evidence. Due to the limited availability of high-quality TAD annotations, supervised deep-learning methods for TAD calling remain rare. To address this bottleneck, we construct improved training annotations by integrating high-coverage Hi-C structure with complementary boundary-associated genomic signals, thereby providing more reliable supervision for model training. We benchmarked ContextTAD against a broad panel of alternative TAD callers across standard comparative evaluation, sequencing-depth robustness analysis, and cross-cell-type transfer settings, and found that it performed strongly against alternative tools across this wide range of settings, with the best overall recovery of biologically supported TADs.

**Availability:** https://github.com/ai4nucleome/ContextTAD

## Introduction

Topologically associating domains (TADs) are a major layer of 3D genome organization and partition chromosomes into local regulatory neighbourhoods (Beagan and Phillips-Cremins, 2020). In Hi-C contact maps, they appear as interaction-enriched blocks near the main diagonal, often with nested substructure at finer scales (Rao et al., 2014). Accurate TAD annotation is therefore important both for describing genome folding and for interpreting boundary-associated architectural factors such as CTCF and cohesin. In practice, however, robust TAD calling remains difficult at 5-kb resolution and typical sequencing depth, where sparse contacts, weak local contrast, and ambiguous nested patterns can obscure biologically meaningful domains.

A broad range of computational methods has been developed for this problem (Zufferey et al., 2018; Sefer, 2022). Early callers relied on one-dimensional summaries of contact asymmetry or insulation (Dixon et al., 2012; Shin et al., 2016), whereas later methods incorporated matrix segmentation, local contrast, and explicit handling of hierarchical domains (Filippova et al., 2014; Rao et al., 2014; An et al., 2019). Learning-based approaches have started to enter this space as well, either by improving boundary annotation directly or by enhancing Hi-C representations before downstream calling (Wang et al., 2024; Fang et al., 2025). Yet the outputs of different callers often remain inconsistent, especially when domain signals are weak or multiple nested structures compete for the same local boundary evidence (Zufferey et al., 2018).

This inconsistency is not only a boundary-detection problem. Many effective TAD callers still follow a boundary-to-domain pipeline (Haddad et al., 2017; Wang et al., 2017; An et al., 2019; Wang et al., 2025), which means that final performance depends on whether predicted boundaries can be assembled into coherent left-right pairs. RobusTAD (Zhang et al., 2025) makes this dependency particularly explicit: it models left and right boundaries separately and then reconstructs nested domains through dynamic programming. deepTAD (Wang et al., 2025) likewise demonstrates that learning can improve boundary annotation, but downstream TAD recovery still depends on how those boundaries are combined. In other words, current pipelines may account for domain compatibility during decoding, yet the representation learned during training is still dominated by local boundary identification rather than by domain-level compatibility.

We developed ContextTAD to move this missing structural context into training. ContextTAD is a deep-learning TAD caller that predicts left- and right-boundary tracks from Hi-C contact maps, but it does so using a representation trained to capture which boundary combinations are compatible with plausible domain structure within a broader local Hi-C window. A second obstacle is that, due to the limited availability of high-quality TAD annotations, supervised deep learning for direct TAD calling remains rare. To address this bottleneck, we construct improved training annotations by integrating high-coverage Hi-C structure with complementary boundary-associated genomic signals, and then use these training annotations as supervision across sequencing depths. With this design, ContextTAD achieves the strongest supported-domain recovery on the GM12878 benchmark, remains robust under progressive downsampling, and transfers effectively to additional cell types.

## Materials and Methods

### ContextTAD Architecture

Figure 1 summarizes the overall study design and the ContextTAD architecture. ContextTAD learns boundary representations for TAD calling under broader local, TAD-scale structural context, rather than predicting intervals directly from raw Hi-C patches. As shown in Fig. 1b, each local O/E Hi-C window is treated as a structured 2D input, but the prediction target remains a pair of 1D boundary tracks aligned to genomic bins. The overall architecture follows three stages: a Hi-C-specific input adaptation stage, a pretrained visual encoding stage, and a boundary-prediction stage that converts 2D context into left- and right-boundary evidence. The predicted boundary tracks are then used for TAD construction from boundary pairs.

**Figure 1.**
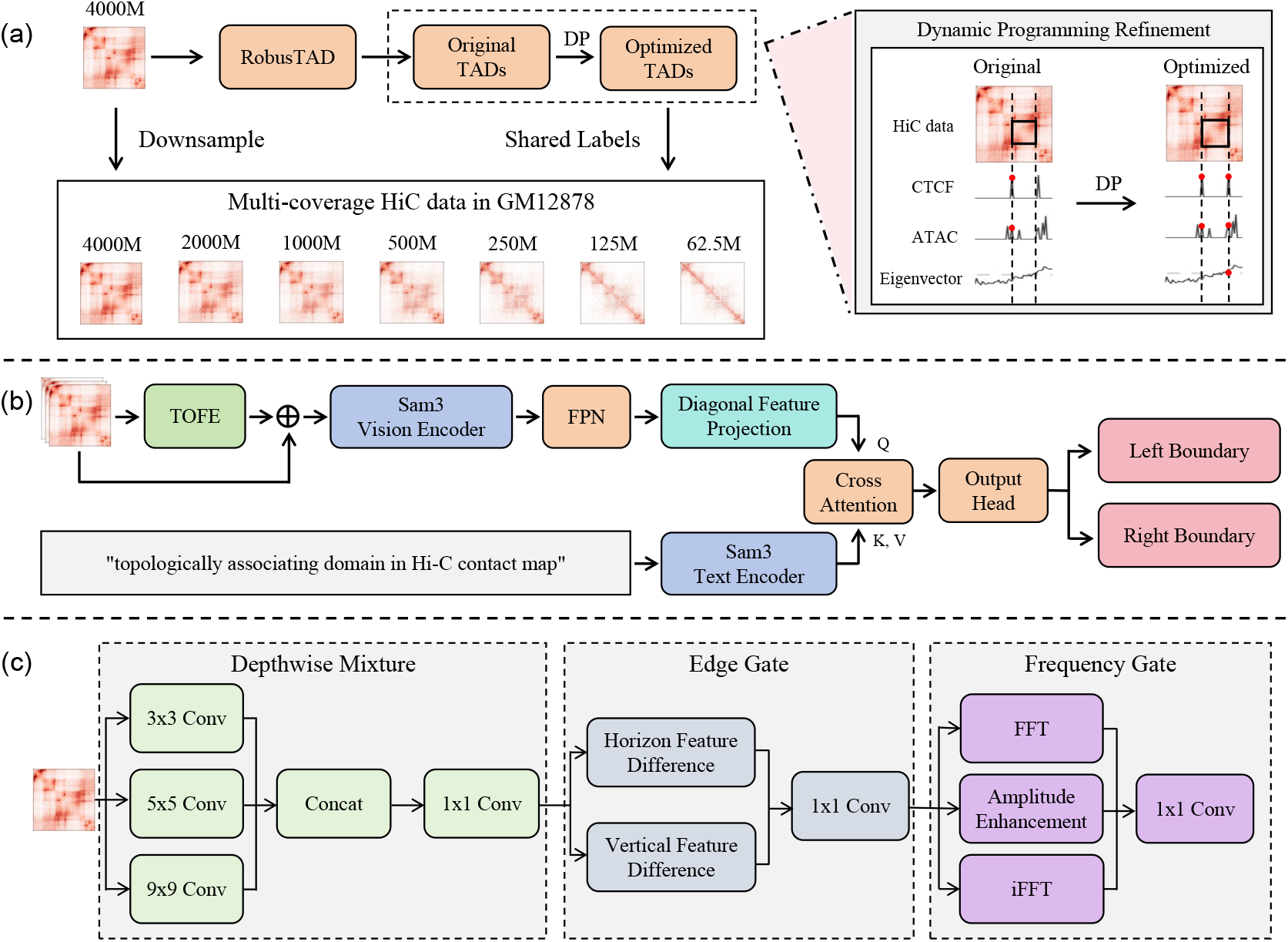
Overview of the study design and the ContextTAD framework. (a) Data preparation and annotation construction from multi-coverage GM12878 Hi-C data. (b) The ContextTAD architecture for context-aware boundary learning from Hi-C contact maps. (c) Internal design of the TOFE module.

To capture 2D interaction patterns from Hi-C windows, we build ContextTAD on top of a pretrained SAM3 (Carion et al., 2026) vision backbone with parameter-efficient adaptation. SAM3 was developed for natural images rather than Hi-C contact maps, so we introduced the TAD O/E Feature Enhancement (TOFE) module as an adaptation layer before visual encoding. As illustrated in Fig. 1c, TOFE processes the input through three branches: the depthwise-mixing branch extracts local domain-related features across multiple spatial scales, the edge gate emphasizes boundary transitions associated with TAD delineation, and the frequency gate modulates characteristic amplitude patterns in Hi-C contact maps. The resulting representation is then processed by the SAM3 vision backbone. Multiscale features are aggregated by a Feature Pyramid Network (FPN) module (Lin et al., 2017), converted into a diagonal-aligned 1D sequence, and combined with a fixed text embedding used as a semantic prior. The final head outputs left-boundary and right-boundary scores for each genomic bin. These boundary tracks are subsequently used for TAD construction. Ablation results for TOFE, the text branch, and the input representation are summarized in Supplementary Note S2.

### Loss Function

ContextTAD produces two boundary probability tracks, one for candidate left endpoints and one for candidate right endpoints. The training objective therefore has two goals. First, the model should identify true TAD boundaries. Second, boundary prediction should reflect the fact that a TAD is defined by a matched left-right boundary pair rather than by isolated boundary peaks. Using paired endpoints as supervision helps the model distinguish boundaries that participate in real domains from local false positives that look boundary-like in isolation. We therefore optimize two complementary terms: a pair loss that rewards endpoint pairs from annotated TADs and suppresses implausible combinations, and a count loss that provides additional supervision to control positive boundary predictions within each window.

#### Pair loss

Let 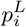 and 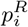 denote the predicted probabilities that bin *i* is a left or right boundary, respectively. For each training window, let 𝒯 denote the set of endpoint pairs defined by the annotated TADs in the training annotations, and let 𝒩 denote a negative set of endpoint pairs that do not correspond to annotated TADs, generated by mismatching true endpoints and by introducing local boundary perturbations. We score a candidate interval (*l, r*) by the product 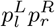, which matches the factorized form later used for TAD construction, and optimize the following objective:

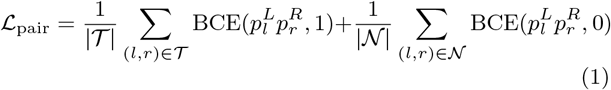

Here, BCE denotes binary cross-entropy. This objective pushes the model to assign high joint support to endpoint pairs that correspond to annotated domains and low support to incompatible pairs. In this way, the two boundary tracks are trained not only to highlight local boundary-like bins, but also to reflect whether those bins can participate in a plausible TAD.

#### Count loss

Boundary prediction in a Hi-C window is a dense prediction problem, and many bins can exhibit weak boundary-like patterns. Pair supervision alone does not fully constrain these diffuse activations. We therefore add a count-aware auxiliary supervision term that provides window-level control over positive boundary prediction:

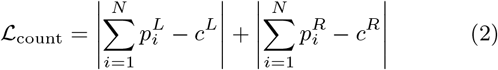

Here, *N* is the number of bins in the current window, and *c*^*L*^ and *c*^*R*^ are the target counts of left and right endpoints derived from the training annotations. These targets are multiplicity-aware: when a shared boundary participates in multiple nested domains, it contributes once for each participation. This auxiliary term stabilizes dense prediction and suppresses spurious positive boundary activations before downstream TAD assembly.

The final training objective is the sum of the two terms:

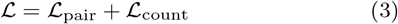

Ablation results for both loss components are summarized in Supplementary Note S2.

### TAD Construction from Boundary Pairs

After inference on overlapping Hi-C windows, the predicted left-boundary and right-boundary scores were aggregated into chromosome-level boundary strength tracks. Candidate TADs were then generated by pairing left and right boundary positions under explicit interval constraints, including valid boundary order and TAD size limits. For each candidate interval (*l, r*), we defined its confidence score as

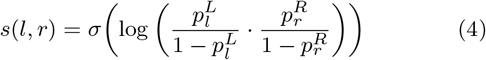

where 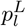 and 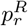 denote the aggregated left-boundary and right-boundary strengths at bins *l* and *r*, respectively, and *σ*(·) is the sigmoid function. This formulation corresponds to a multiplicative fusion of left-boundary and right-boundary evidence in odds space, so that a candidate receives a high score only when both endpoints are strongly supported.

The candidates were then consolidated by post-processing. We first removed intervals that were shorter than the minimum allowed TAD size (5 bins) or longer than the maximum allowed size (350 bins). We next applied overlap-based deduplication to suppress near-duplicate candidates, retaining the higher-confidence interval when strongly overlapping candidates conflicted. Finally, nearby candidates (within 2 bins) with highly similar boundaries were merged by boundary snapping to reduce redundancy caused by local score fluctuations. The final TAD set for each chromosome was obtained from the remaining non-redundant candidate intervals.

### Training Data Construction

#### Data preprocessing

As summarized in Fig. 1a, we created a series of downsampled GM12878 Hi-C datasets spanning seven sequencing-depth settings, from 4000M to 62.5M valid contact pairs. All analyses were performed at 5-kb resolution. Balanced contacts were converted into observed-over-expected (O/E) matrices and partitioned into overlapping windows of 400 bins with a stride of 200 bins. Supervision was constructed once from the highest-coverage 4000M map and then reused for the corresponding genomic windows across all downsampled coverages, so that the learning problem isolates changes in Hi-C signal quality rather than changes in TAD annotation definition. We used chromosomes 11 and 12 for validation, chromosomes 15-17 for testing, and the remaining autosomes for training. K562 and IMR90 were processed with the same windowing scheme and reserved for external evaluation. Dataset statistics are summarized in Table 1.

**Table 1.** Dataset statistics.

| Cell line | GM12878 |  |  | K562 | IMR90 |
| --- | --- | --- | --- | --- | --- |
| Split | Train | Val | Test | Test | Test |
| Coverage | 62.5M, 125M, 250M, 500M, 1000M, 2000M, 4000M |  |  | 300M | 1000M |
| Chromosomes | Remaining autosomes | 11, 12 | 15, 16, 17 | 15, 16, 17 | 15, 16, 17 |
| Num. windows | 2213 | 265 | 242 | 274 | 274 |

#### Dynamic programming-based annotation construction

Due to the limited availability of high-quality TAD annotations, supervised deep learning for TAD calling remains rare. We therefore introduced an annotation-construction stage instead of directly training on raw outputs from an existing caller. We started from the initial RobusTAD annotations on the 4000M GM12878 map and refined them by dynamic programming to reposition local boundaries while preserving the overall domain layout. Refinement was performed jointly for nearby or nested domains within each local component so that shared boundaries were optimized together. The optimization objective was to locally adjust TAD boundaries so that the resulting domains maximized agreement with local Hi-C structural features and boundary-associated genomic signals, including CTCF occupancy and TAD corner patterns. For each component 𝒞, candidate boundary positions were restricted to a local neighborhood around the initial boundaries, and the optimal boundary assignment was obtained by

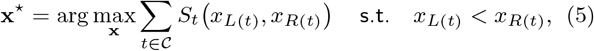

where *L*(*t*) and *R*(*t*) denote the left and right boundaries of a TAD, respectively. The single-domain score was defined as

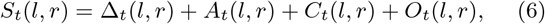

Here, Δ_*t*_(*l, r*) is the RobusTAD structural score, *A*_*t*_(*l, r*) measures alignment with orthogonal boundary-associated genomic signals, *C*_*t*_(*l, r*) evaluates local Hi-C corner consistency, and *O*_*t*_(*l, r*) is a locality term favoring minimal displacement from the initial annotation. The genomic signals used in *A*_*t*_ included CTCF ChIP-seq occupancy, local chromatin accessibility measured by ATAC-seq, and compartment-transition signatures captured by eigenvector tracks (Rao et al., 2014; Lieberman-Aiden et al., 2009; Pletenev et al., 2024). The resulting 4000M training annotations were then fixed and reused as supervision across all sequencing-depth settings. Additional scoring details and search procedures are described in Supplementary Note S1.

### Training strategy

We adopted a random-single coverage sampling strategy as each genomic window was available at seven sequencing depths but shared the same training annotation. At each epoch, each genomic window contributed one randomly selected coverage version for training, and validation followed the same protocol. In this way, ContextTAD was repeatedly exposed to Hi-C inputs of varying signal quality while maintaining a unified supervision target.

During optimization, the backbones from SAM3 were kept largely frozen, and only the vision branch (Bolya et al., 2025) was adapted through LoRA (Hu et al., 2022), whereas the text branch remained frozen. The newly introduced modules, including TOFE, FPN, diagonal projection, and the boundary prediction head, were trained from scratch. We optimized the model with AdamW (Loshchilov and Hutter, 2019) using a learning rate of 3 × 10^−4^, weight decay of 0.01, LoRA rank of 16, and cosine learning-rate decay with warmup. All experiments were run with a fixed random seed of 42, and the model was trained for 5 epochs.

### Evaluation Protocol

We evaluated ContextTAD under the GM12878 benchmark setting at 5-kb resolution. The main comparison was performed on the held-out test chromosomes chr15, chr16, and chr17 using the 250M GM12878 Hi-C map, following prior TAD-calling benchmarks (Zufferey et al., 2018; Zhang et al., 2025). We further applied the same protocol across all seven GM12878 coverage levels to assess sequencing-depth robustness, and to K562 and IMR90 to assess cross-cell-type transfer. Baseline comparisons included representative classical, hierarchical, and learning-based TAD callers.

The primary evaluation metric was the number of CTCF ChIA-PET-supported TADs. Follow RobusTAD, we defined TADs that do not contain any smaller TADs as level 0 TADs, and TADs that contain at least one TADs as level 1+ TADs. We also evaluated boundary support using strand-matched CTCF ChIP-seq peaks and measured enrichment of CTCF, RAD21, and SMC3 around predicted boundaries. For sequencing-depth experiments, we additionally reported the total number of predicted TADs together with the number of supported TADs. The same domain-level and boundary-level evaluation protocol was used for K562 and IMR90.

## Results

### ContextTAD performs strongly on a typical-coverage Hi-C map

On the GM12878 250M benchmark, which represents a typical-coverage Hi-C setting, we compared ContextTAD against 15 existing TAD callers using the biological support metrics summarized in Fig. 2: Grinch (Lee and Roy, 2021), CaTCH (Zhan et al., 2017), Domaincall (Dixon et al., 2012), Arrowhead (Rao et al., 2014), deDoc (Li et al., 2018), EAST (Roayaei Ardakany and Lonardi, 2017), HiCSeg (Lévy-Leduc et al., 2014), GMAP (Yu et al., 2017), HiTAD (Wang et al., 2017), IC-Finder (Haddad et al., 2017), TopDom (Shin et al., 2016), Armatus (Filippova et al., 2014), OnTAD (An et al., 2019), RefHiC (Zhang and Blanchette, 2022), and RobusTAD (Zhang et al., 2025). ContextTAD achieved the highest number of CTCF ChIA-PET-supported TADs among all compared methods (Fig. 2a). This advantage was evident for both level-0 domains and nested level-1+ domains, indicating that the gain was not restricted to one specific structural regime. RobusTAD and RefHiC formed the strongest competing group, whereas Arrowhead behaved as a conservative caller with relatively few but often highly supported predictions. OnTAD and Armatus also detected nested TADs, although the fraction of ChIA-PET-supported TADs was slightly lower than that of ContextTAD. Overall, ContextTAD occupied the most favorable yield-versus-support operating point in this benchmark.

**Figure 2.**
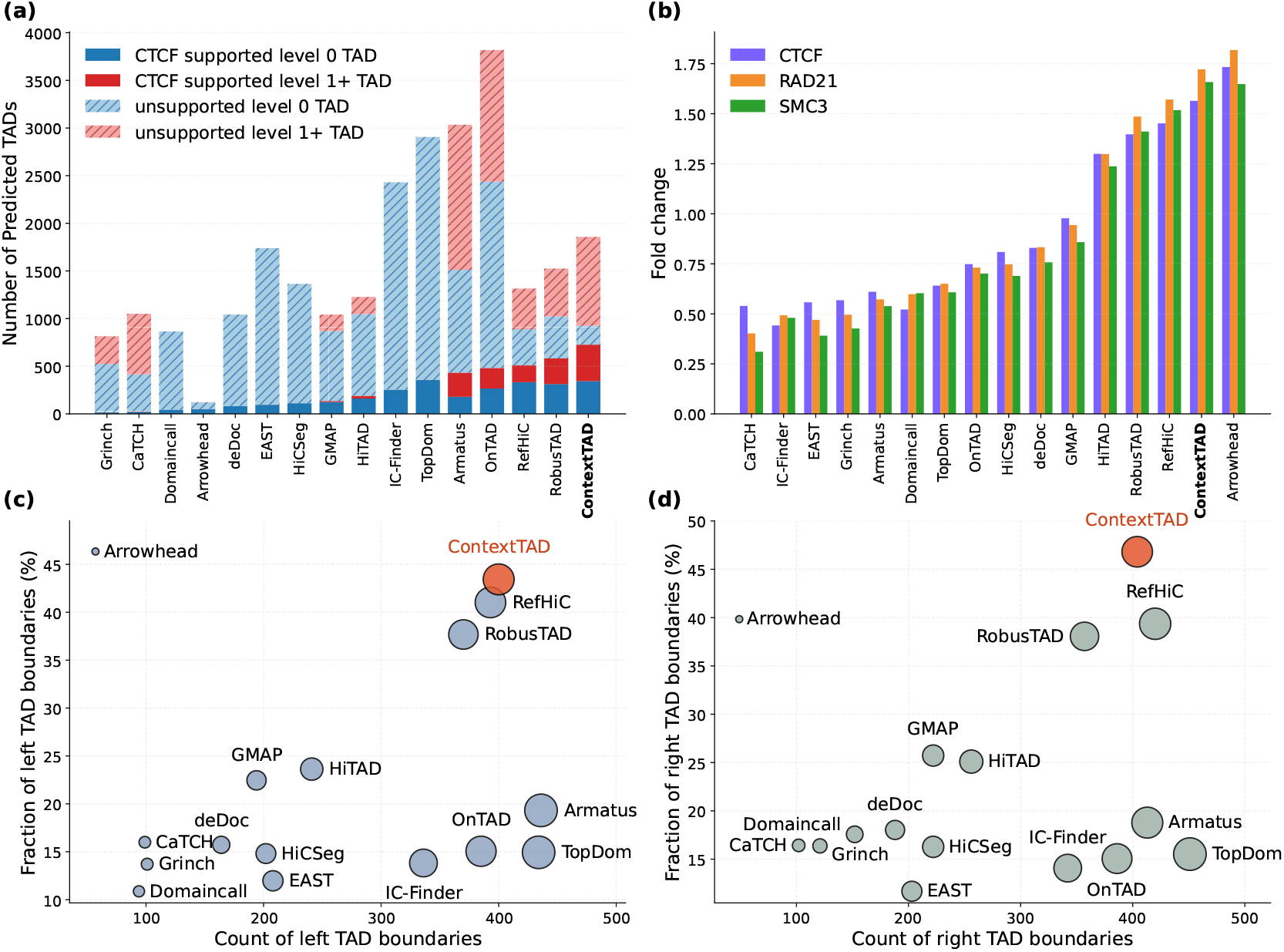
Comparison of ContextTAD and existing TAD callers on GM12878 Hi-C data at 250M valid contact pairs. (a) CTCF ChIA-PET-supported TAD count stratified by nesting level. (b) Fold change of architectural proteins around predicted TAD boundaries. (c) Left-boundary support evaluated by CTCF ChIP-seq peaks. (d) Right-boundary support evaluated by CTCF ChIP-seq peaks.

This gain did not come from sacrificing boundary quality. ContextTAD remained within the top-performing group for both the number and fraction of strand-supported left and right boundaries (Fig. 2c,d), showing that the learned boundary tracks remained biologically consistent at the endpoint level. ContextTAD also achieved strong enrichment of canonical architectural proteins, including CTCF, RAD21, and SMC3, around predicted boundaries (Fig. 2b). Although a few more conservative callers obtained slightly higher enrichment for individual proteins, those gains were typically accompanied by much smaller TAD sets. ContextTAD therefore improved supported-domain recovery while maintaining strong boundary support and architectural-protein enrichment. Aggregate pileup visualizations comparing ContextTAD with all callers are shown in Supplementary Fig. S4, and representative examples are shown in Supplementary Fig. S5.

### ContextTAD maintains strong TAD recovery across sequencing depths

We next evaluated whether ContextTAD remained effective as sequencing depth decreased. Across the seven downsampled GM12878 datasets, ContextTAD maintained high recovery of CTCF-supported TADs over a broad coverage range (Fig. 3a,b). RobusTAD recovered slightly more supported TADs at 4000M, but its supported TAD count decreased more rapidly as coverage declined. From 2000M downward, ContextTAD recovered the largest number of supported TADs among the compared methods. RefHiC followed a relatively stable pattern, although both its total TAD count and its supported TAD count remained below those of ContextTAD. OnTAD continued to report many candidate domains at lower coverage, but the number of supported TADs remained lower than that of ContextTAD. These results show that ContextTAD preserves biologically supported TAD recovery when Hi-C signal becomes sparse.

**Figure 3.**
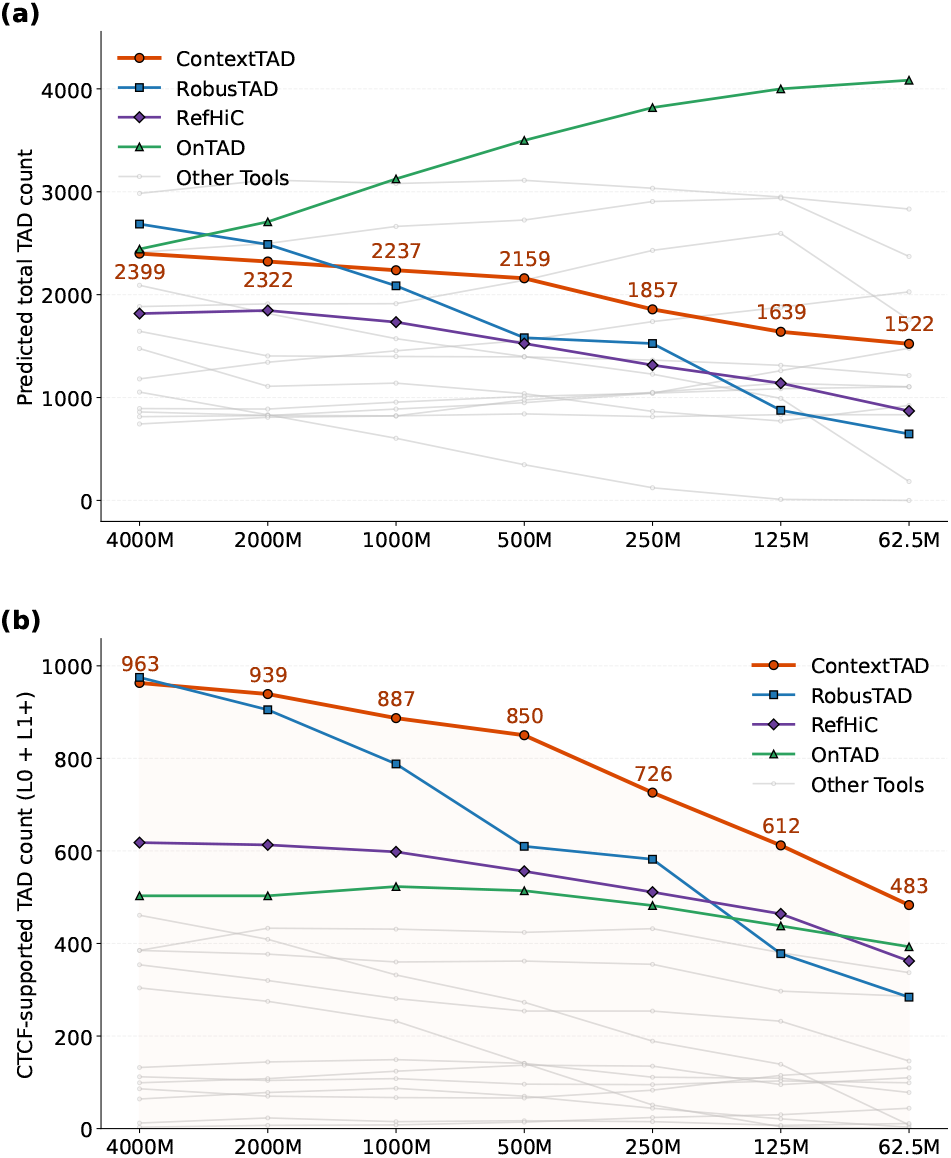
Comparison of robustness for ContextTAD and 15 tools in downsampled GM12878 Hi-C data. (a) Total number of predicted TADs across coverages. (b) Number of TADs supported by CTCF ChIA-PET data across coverages.

### ContextTAD transfers across cell types without retraining

We then tested whether a model trained on GM12878 could be applied directly to other cell types. As shown in Fig. 4a,b, ContextTAD retained strong domain-level performance without retraining. In K562, which represents the lower-coverage external setting at 300M, ContextTAD recovered the largest number of CTCF-supported TADs among the compared methods. In IMR90, ContextTAD remained competitive with the strongest baselines, although RobusTAD and RefHiC recovered slightly more supported TADs. These results indicate that the boundary representation learned by ContextTAD generalizes beyond the training cell line.

**Figure 4.**
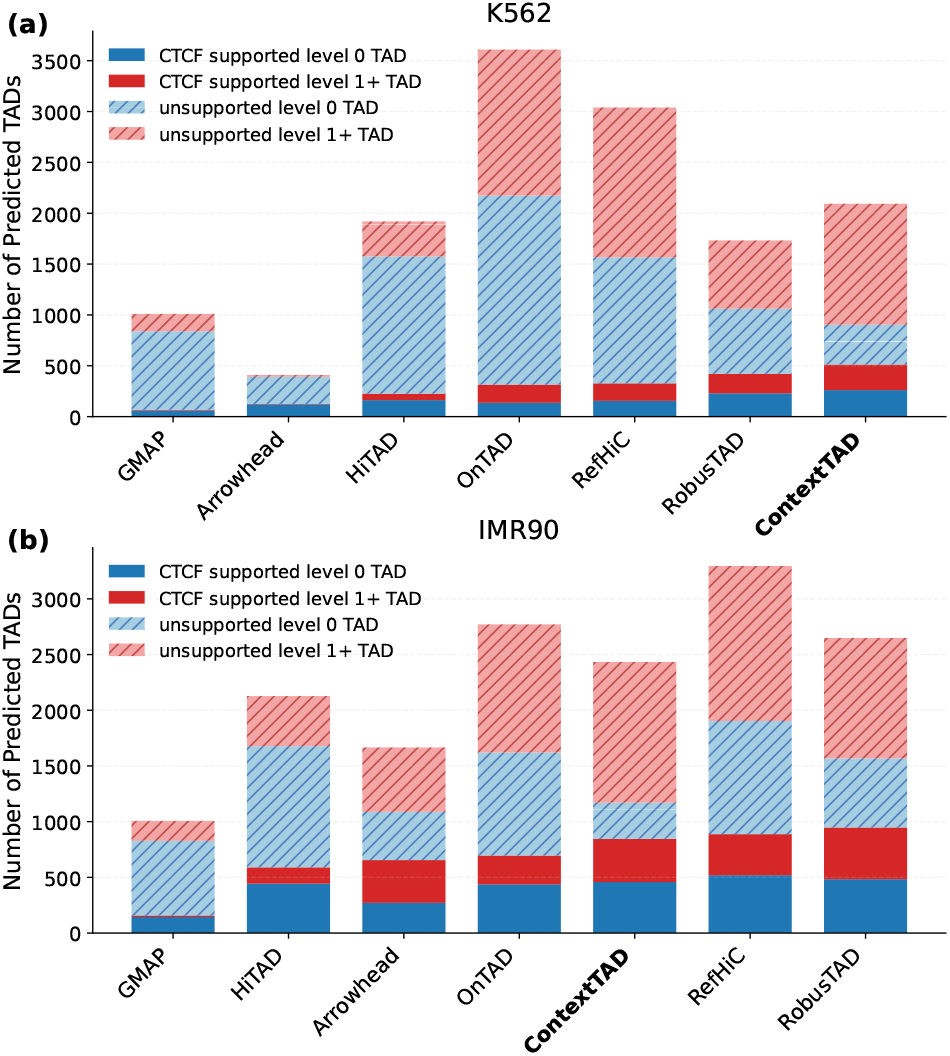
Comparison of ContextTAD and other six TAD callers on Hi-C data derived from IMR-90 and K562 cell lines. (a) and (b) CTCF ChIA-PET-supported TAD count stratified by nesting level.

Boundary-centered CTCF profiles in Supplementary Fig. S3 support the same conclusion. In both K562 and IMR90, forward-strand CTCF signal around predicted left boundaries and reverse-strand CTCF signal around predicted right boundaries remained centered near the predicted boundary position, indicating that the directional boundary pattern learned from GM12878 was preserved after transfer.

## Discussion and Conclusion

In this paper, we presented ContextTAD, a context-aware deep-learning framework for TAD calling from Hi-C contact maps. ContextTAD learns boundary evidence from broader local Hi-C context and combines compatibility-aware supervision with improved training-data construction to make supervised TAD calling practical. Across typical-coverage, low-coverage, and cross-cell-type evaluation settings, ContextTAD consistently recovered large numbers of biologically supported TADs while maintaining strong boundary-level biological consistency. These results support context-aware boundary learning as a practical and effective direction for supervised TAD calling.

For boundary-to-domain TAD calling pipelines, the quality of the learned boundary evidence is as important as the downstream TAD construction procedure. In many existing pipelines, domain compatibility is considered mainly after boundary prediction, whereas ContextTAD moves part of this structural information into training through broader local Hi-C context and compatibility-aware supervision. Our results suggest that this design is particularly useful when Hi-C signal is weak, because the advantage of ContextTAD was preserved under progressive downsampling and remained visible after transfer to additional cell types.

Supervised TAD learning is limited not only by model architecture, but also by the availability of reliable training annotations. In this study, we addressed this bottleneck by constructing training annotations from high-coverage Hi-C maps together with complementary boundary-associated genomic signals. This annotation-construction step made it possible to train a supervised model using a more stable domain definition than would be obtained by directly inheriting raw outputs from an existing caller. This strategy may also be useful for other supervised learning tasks in Hi-C data analysis.

Despite its strong performance across typical-coverage, low-coverage, and cross-cell-type evaluation settings, ContextTAD still constructs domains after boundary prediction rather than predicting TADs directly, and therefore cannot fully benefit from end-to-end deep learning for TAD calling. We expect that future end-to-end frameworks for direct TAD prediction will allow deep learning to be more fully exploited for this task.

## Supporting information

All supplemental files

## Author contributions

W.L. and Y.Z. conceived the study. W.L. and Y.H. performed the analysis. Y.Z. supervised the project. W.L. and Y.Z. wrote the manuscript. All authors read and approved the final version.

## Acknowledgements

This work was supported by the National Natural Science Foundation of China (No. 32500550). Y.Z. was supported by the Ministry of Human Resources and Social Security of the People’s Republic of China (No. Y20250128) and a Guangdong Provincial Project (2024QN11N085).

## Data and code availability

The datasets and codebase used in this work can be accessed at https://github.com/ai4nucleome/ContextTAD.

