## Supplementary material for "ContextTAD: Context-aware boundary learning for TAD calling from Hi-C contact maps": All supplemental files

### Supplementary Note S1. Details of dynamic programming-based annotation refinement

This note describes how we refined the initial RobusTAD annotations on the 4000M GM12878 map by dynamic programming. The goal was to locally adjust TAD boundaries while preserving the overall domain layout. Throughout this procedure, the input TAD annotations were stored in half-open coordinates $[l, r)$, whereas internal optimization used the corresponding closed interval $[l, r-1]$ so that corner consistency and boundary-alignment checks were evaluated on the actual terminal bins. After optimization, the refined annotations were converted back to half-open coordinates for downstream use.

**Boundary clustering and candidate generation.** Nearby boundaries from the initial RobusTAD annotations were first clustered into unique boundaries so that shared or nearly shared boundaries could be optimized jointly. Specifically, two starting boundaries were assigned to the same unique boundary if their coordinates differed by at most 3 bins. For each unique boundary $b$ with reference coordinate $\hat{b}$, we defined a local candidate set

$$\mathcal{P}_b = \{\max(0, \hat{b} - 5), \ldots, \min(n - 1, \hat{b} + 5)\}, \tag{1}$$

where $n$ is the number of bins in the current window. Candidate positions were therefore restricted to a local neighborhood around the starting annotation rather than searched globally.

**Connected components of coupled TADs.** Refinement was performed jointly for TADs that shared the same unique left boundary or the same unique right boundary. We identified such groups with a union-find data structure. Importantly, only boundaries of the same type were considered connected. Thus, if the right boundary of one TAD coincided with the left boundary of another, the two TADs were not merged solely because of this end-start adjacency. This design allowed nearby or nested domains with truly shared boundaries to be optimized coherently while avoiding spurious coupling between adjacent but independent domains.

**Joint optimization within each component.** Let $\mathcal{C}$ denote a connected component of coupled TADs, and let $\mathcal{B}(\mathcal{C})$ be the set of unique boundaries that appear in this component. For each TAD $t \in \mathcal{C}$, let $L(t)$ and $R(t)$ denote the unique left and right boundaries associated with that TAD. The refined boundary configuration was obtained by

$$\mathbf{x}^\star = \arg \max_{\mathbf{x} \in \prod_{b \in \mathcal{B}(\mathcal{C})} \mathcal{P}_b} \sum_{t \in \mathcal{C}} S_t\big(x_{L(t)}, x_{R(t)}\big) \quad \text{s.t.} \quad x_{L(t)} < x_{R(t)} \;\; \forall t \in \mathcal{C}, \tag{2}$$

where $\mathbf{x}$ is a joint assignment of candidate coordinates to all unique boundaries in the component. We used exhaustive search when the total number of candidate combinations did not exceed $10^5$. For larger components, we used a coordinate-wise iterative approximation, in which each boundary was updated by scanning its candidate set while fixing the other boundaries in the component.

**Single-domain score.** The score of a candidate TAD $t$ under boundary assignment $(l, r)$ was defined as

$$S_t(l, r) = \Delta_t(l, r) + w_t A_t(l, r) + C_t(l, r) + Q_t(l, r) + O_t(l, r), \tag{3}$$

where $\Delta_t(l, r)$ is the RobusTAD structural score, $A_t(l, r)$ is the genomic-signal alignment term, $C_t(l, r)$ is the local corner-consistency term, $Q_t(l, r)$ is a local score-optimality term, and $O_t(l, r)$ is a displacement term that discourages unnecessary movement away from the starting annotation. The weight

$$w_t = \max\left(1, \frac{s_{L(t)} + s_{R(t)}}{2}\right) \tag{4}$$

upweights alignment rewards when the left or right boundary is shared by multiple TADs, where $s_{L(t)}$ and $s_{R(t)}$ denote the sharing counts of the corresponding unique boundaries.

**RobusTAD structural term.** The term $\Delta_t(l, r)$ was computed from the RobusTAD scoring function on the larger local Hi-C matrix. Internally, the closed interval $[l, r]$ was converted back to the half-open interval $[l, r + 1)$ before calling the RobusTAD score. This term measures whether within-domain contacts are enriched relative to local flanking contacts and serves as the main structural component of the DP objective.

**Alignment term.** The alignment term was based on three orthogonal one-dimensional signals associated with TAD boundaries: CTCF ChIP-seq occupancy, ATAC-seq accessibility, and eigenvector transitions. We used a priority-based reward scheme,

$$A_t(l,r) = \begin{cases} 100, & \text{both boundaries overlap CTCF peaks,} \\ 80, & \text{one boundary overlaps CTCF and the other is at an eigenvector transition,} \\ 50, & \text{one boundary overlaps CTCF and the other overlaps ATAC,} \\ 30, & \text{both boundaries overlap ATAC peaks,} \\ 20, & \text{only one boundary overlaps CTCF,} \\ 10, & \text{only one boundary overlaps ATAC,} \\ 0, & \text{otherwise.} \end{cases} \tag{5}$$

An eigenvector transition was defined by either a sign change between consecutive bins or an eigenvector magnitude close to zero. These rewards favored boundary placements that were consistent with strong architectural evidence while still allowing joint optimization of shared boundaries.

**Corner-consistency term.** TAD corners in Hi-C maps often coincide with local maxima. We therefore evaluated whether the candidate corner $(l, r)$ was supported by the O/E matrix in a local $3 \times 3$ neighborhood:

$$C_t(l,r) = \begin{cases} 30, & (l,r) \text{ is the maximum in the local } 3 \times 3 \text{ neighborhood,} \\ 15, & \text{the maximum lies at an immediately adjacent lower-left position,} \\ 0, & \text{otherwise.} \end{cases} \tag{6}$$

This term rewards corner-like geometry while remaining tolerant to slight local offsets.

**Local score-optimality term.** To avoid selecting boundary placements with poor structural support, we compared the RobusTAD score of the current candidate to the best score obtained by jointly shifting both boundaries within the same local search window:

$$\rho_t(l,r) = \frac{\Delta_t(l,r)}{\max_{d \in [-5,5]} \Delta_t(l+d, r+d)}. \tag{7}$$

The score-optimality term was then defined as

$$Q_t(l,r) = \begin{cases} 50, & \rho_t(l,r) \geq 0.95, \\ 15, & 0.85 \leq \rho_t(l,r) < 0.95, \\ 0, & 0.80 \leq \rho_t(l,r) < 0.85, \\ -20, & \rho_t(l,r) < 0.80. \end{cases} \tag{8}$$

This soft constraint favors boundary placements that remain close to the best-scoring structural configuration within the local search window without enforcing a hard optimum.

**Displacement term.** Finally, we included a small displacement term that encourages the optimizer not to move boundaries unless there is sufficient evidence to do so:

$$O_t(l, r) = 2.5\,\mathbf{1}[l = l_t^0] + 2.5\,\mathbf{1}[r = r_t^0], \tag{9}$$

where $(l_t^0, r_t^0)$ denotes the initial RobusTAD boundary pair for TAD $t$.

**Output of refinement.** The optimized boundary configuration was written back to the initial TAD list to obtain the refined 4000M annotation for each window. These refined annotations were then fixed and reused as supervision for the same genomic windows across all sequencing-depth settings. As shown in Supplementary Fig. S1 and Supplementary Fig. S2, this refinement improved alignment with boundary-associated genomic signals and local Hi-C corner structure while preserving the original nested domain organization.

### Supplementary Note S2. Ablation study

Supplementary Table S1 summarizes the main ablation results. At the module level, the full ContextTAD model consistently achieved the strongest overall performance. Removing TOFE reduced CTCF-supported TAD recovery across all sequencing depths, indicating that adapting the Hi-C input before visual encoding is important for robust structural feature extraction. Replacing the O/E matrix with the observed matrix also degraded both boundary support and supported-domain recovery, consistent with the idea that O/E normalization better highlights domain-relevant contrast. Removing the text branch led to a smaller drop at high coverage but a clearer decline at lower coverage, suggesting that this branch provides a weak but useful semantic prior, especially when Hi-C signal becomes sparse.

At the objective level, both the pair loss and the count loss were important. Removing the pair loss reduced supported TAD recovery to nearly zero across all coverage settings. This result indicates that local boundary activation alone is insufficient, and that explicit supervision of valid left–right boundary pairing is needed to produce boundary scores that remain useful for TAD construction. Removing the count loss also markedly reduced performance, leaving only a small number of supported domains together with distorted boundary support. These results show that ContextTAD depends not only on the visual backbone and input design, but also on compatibility-aware and count-aware supervision to produce boundary tracks that are suitable for downstream TAD calling.

To further test whether explicit boundary supervision could replace pairwise supervision, we replaced the pair loss with a boundary loss while retaining the count loss. For an input window with $N$ bins, let $p_i^L, p_i^R \in (0,1)$ denote the predicted probabilities that bin $i$ is a left or right boundary, and let $y_i^L, y_i^R \in \{0,1\}$ denote the corresponding ground-truth boundary masks. The boundary loss was defined as

$$\mathcal{L}_{\text{boundary}} = \frac{1}{2}\left(\mathcal{L}_{\text{BCE}}^L + \mathcal{L}_{\text{BCE}}^R\right) + \frac{1}{2}\left(\mathcal{L}_{\text{Dice}}^L + \mathcal{L}_{\text{Dice}}^R\right) + 0.05\left(\mathcal{L}_{\text{smooth}}^L + \mathcal{L}_{\text{smooth}}^R\right). \tag{10}$$

The weighted binary cross-entropy term for the left boundary map was

$$\mathcal{L}_{\text{BCE}}^L = -\frac{1}{N}\sum_{i=1}^{N} w_i^L \left[y_i^L \log p_i^L + (1-y_i^L)\log(1-p_i^L)\right], \tag{11}$$

where

$$w_i^L = \begin{cases} \alpha, & y_i^L = 1, \\ 1, & y_i^L = 0, \end{cases} \tag{12}$$

and $\alpha$ is the positive-class weight. The right-boundary term $\mathcal{L}_{\text{BCE}}^R$ was defined analogously.

The Dice term for the left boundary map was

$$\mathcal{L}_{\text{Dice}}^L = 1 - \frac{2\sum_{i=1}^{N} p_i^L y_i^L + 1}{\sum_{i=1}^{N} p_i^L + \sum_{i=1}^{N} y_i^L + 1}, \tag{13}$$

with $\mathcal{L}_{\text{Dice}}^R$ defined in the same way for the right boundary map.

To encourage local smoothness of the predicted boundary tracks, we used

$$\mathcal{L}_{\text{smooth}}^L = \frac{1}{N-1}\sum_{i=1}^{N-1}\left|p_{i+1}^L - p_i^L\right|, \qquad \mathcal{L}_{\text{smooth}}^R = \frac{1}{N-1}\sum_{i=1}^{N-1}\left|p_{i+1}^R - p_i^R\right|. \tag{14}$$

Despite this explicit marginal supervision on left and right boundary maps, replacing the pair loss with $\mathcal{L}_{\text{boundary}}$ did not restore performance. Even with the count loss retained, supported TAD recovery remained zero across all coverage settings (Supplementary Table S1). At 250M coverage, only weak boundary support remained (17 CTCF-supported left boundaries and 22 CTCF-supported right boundaries). These results indicate that marginal boundary supervision alone cannot replace explicit pairwise endpoint coupling under our factorized TAD-construction strategy.

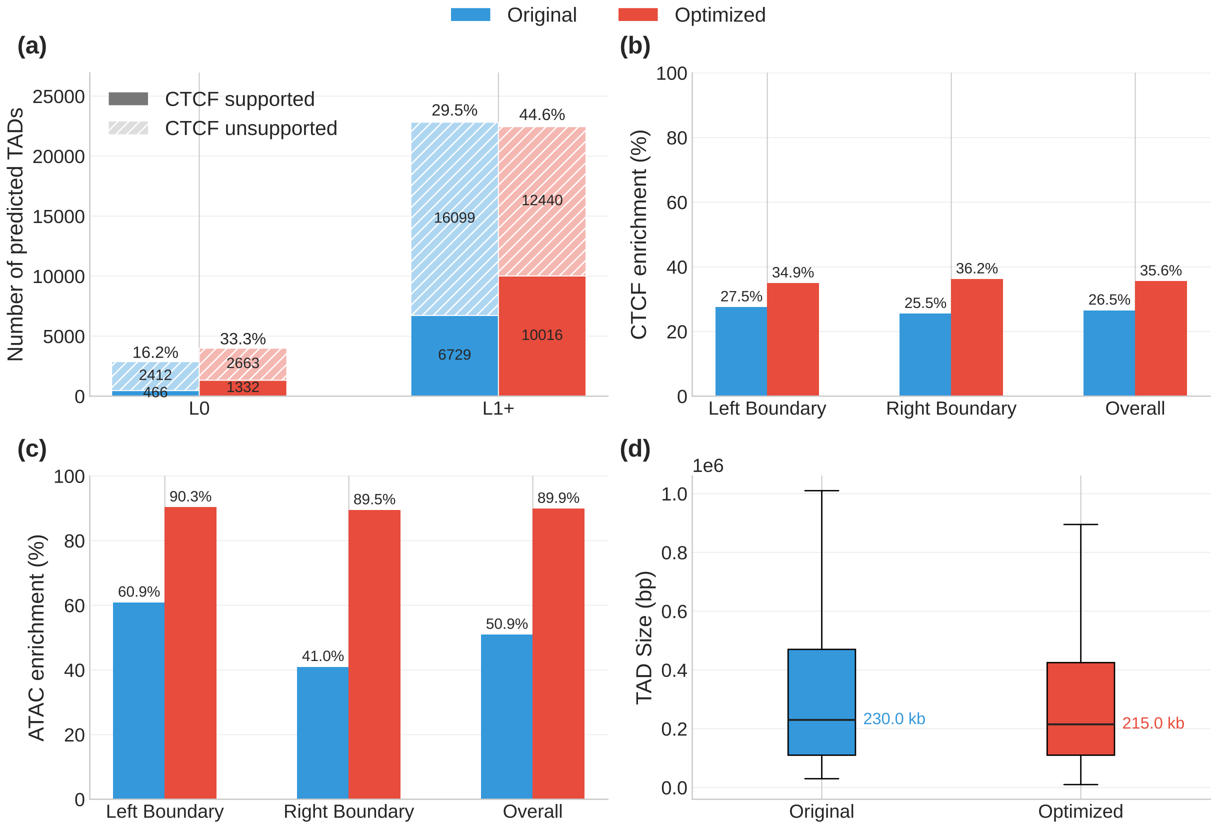


Supplementary Figure S1. Effect of dynamic-programming annotation refinement on the initial TAD annotations (all autosomes). (A) Numbers and fractions of CTCF ChIA-PET supported TADs before and after refinement, stratified by nesting level. (B) Enrichment at left, right, and overall boundaries supported by CTCF ChIP-seq data. (C) Enrichment at left, right, and overall boundaries supported by ATAC-seq data. (D) Distribution of TAD sizes before and after refinement. Dynamic-programming refinement increases biological support and boundary-associated signal enrichment while preserving the overall TAD size distribution.

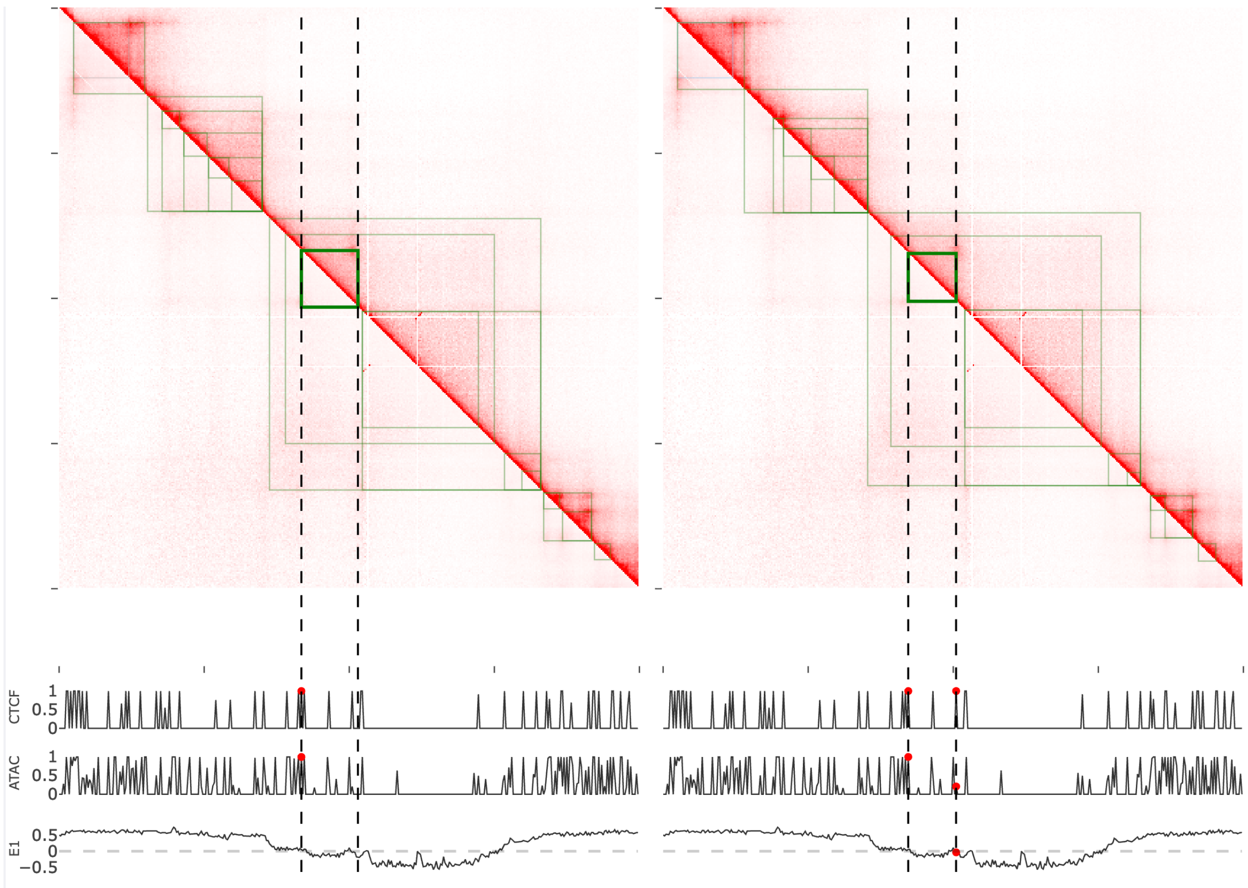


Supplementary Figure S2. An example of local boundary refinement by dynamic programming. Left, initial TAD annotations. Right, optimized annotations after refinement. The dashed lines indicate the local boundary region under refinement, and the highlighted domain shows how the refined boundary placement better aligns with the local Hi-C corner pattern and supporting one-dimensional genomic signals, including CTCF ChIP-seq, ATAC-seq, and eigenvector transitions.

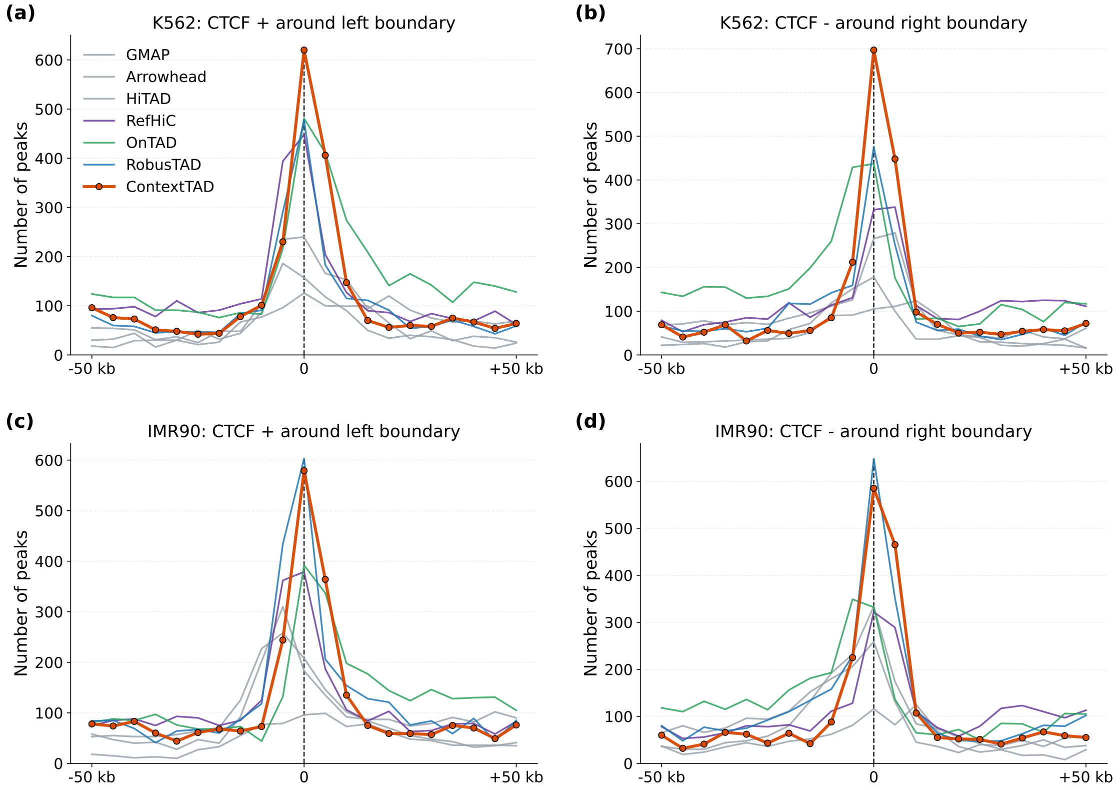


Supplementary Figure S3. Boundary-centered CTCF profiles in external cell types. (a),(b) Forward- and reverse-strand CTCF occupancy around predicted left and right boundaries in K562. (c),(d) The corresponding profiles in IMR90. ContextTAD retains sharply centered directional CTCF enrichment relative to representative baselines in both transfer settings, supporting the portability of the learned boundary representation

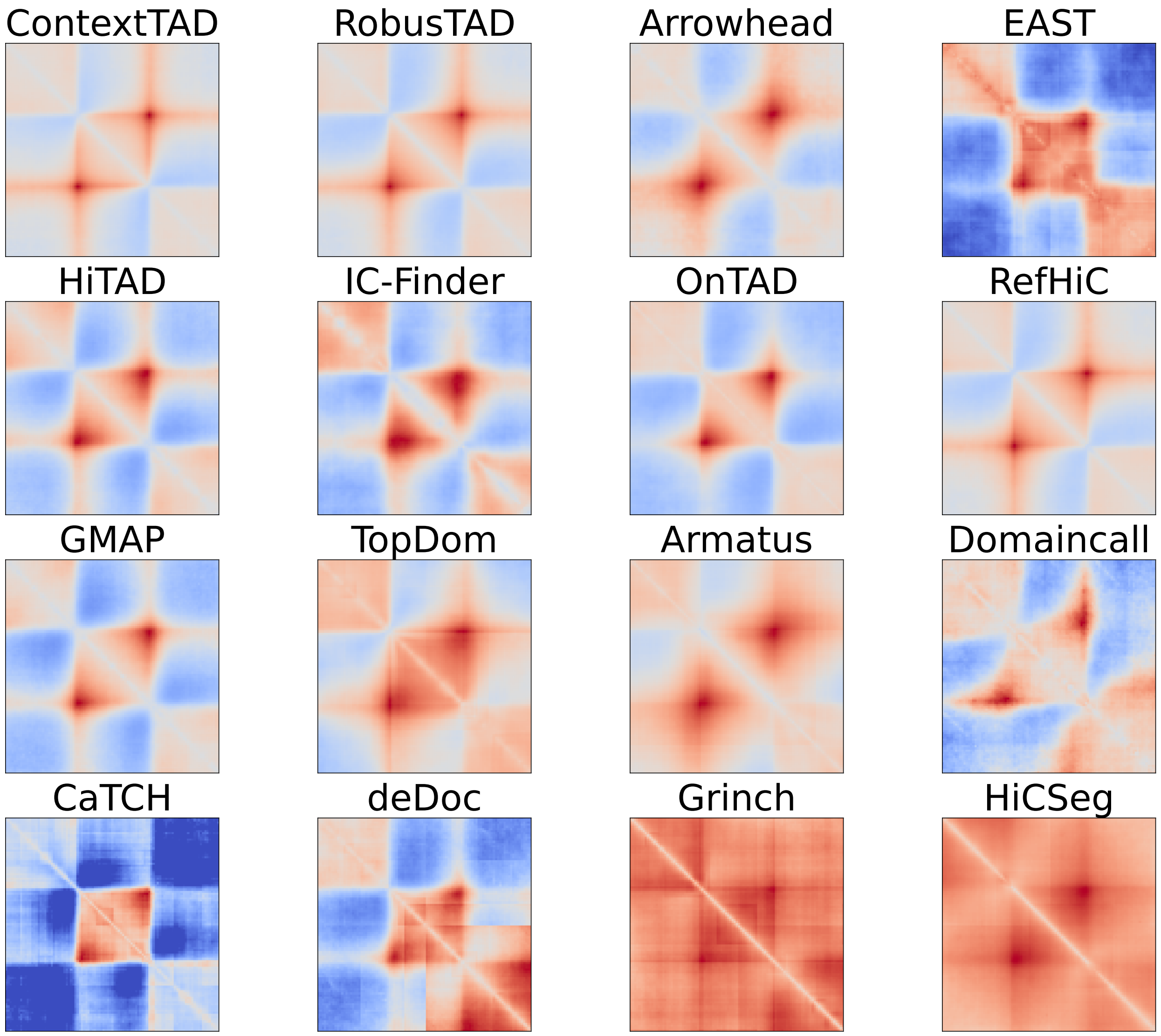


Supplementary Figure S4. Visual comparison of TADs predicted by ContextTAD and 15 existing callers from GM12878 Hi-C data. These rescaled pileup plots were created by aggregating the TAD predictions used in the GM12878 benchmark at 250M coverage. ContextTAD, RobusTAD, RefHiC, and a small number of other strong callers recover canonical TAD corner structure, whereas weaker callers produce blurrier or less coherent aggregate patterns.

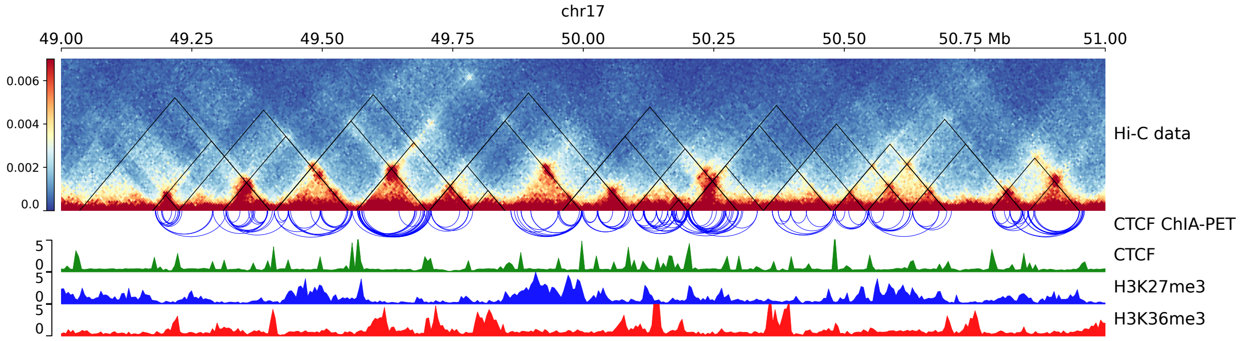


Supplementary Figure S5. ContextTAD predictions for a representative genomic region on chr17 in GM12878 cells. Predicted TADs are overlaid on the local Hi-C contact map together with CTCF ChIA-PET, CTCF ChIP-seq, H3K27me3, and H3K36me3 tracks. ContextTAD predictions align with local interaction blocks and are supported by multiple CTCF-mediated chromatin loops and boundary-associated CTCF enrichment, while neighboring domains exhibit distinct chromatin-state patterns.

| Model | CTCF ChIP-seq supported count(ratio), 250M | | CTCF ChIA-PET supported count | | | | | | |
|---|---|---|---|---|---|---|---|---|---|
| | Left | Right | 4000M | 2000M | 1000M | 500M | 250M | 125M | 62.5M |
| **Baseline** | **400 (43.43%)** | **404 (46.81%)** | **963** | **939** | **887** | **850** | **726** | **612** | **483** |
| Module Ablation | | | | | | | | | |
| No TOFE | 385(42.73%) | 395(48.70%) | 890 | 862 | 826 | 798 | 652 | 582 | 457 |
| No Text Branch | 393 (44.31%) | 355 (48.10%) | 915 | 894 | 828 | 787 | 618 | 508 | 340 |
| Observed Matrix | 385 (39.69%) | 378 (42.14%) | 935 | 925 | 880 | 830 | 671 | 562 | 440 |
| Loss Ablation | | | | | | | | | |
| No Pair loss | 15 (9.87%) | 15 (10.00%) | 0 | 0 | 0 | 0 | 0 | 0 | 0 |
| No Count loss | 30 (8.26%) | 56 (70%) | 76 | 74 | 68 | 70 | 74 | 68 | 65 |
| Boundary + Count loss | 17 (10.97%) | 22 (14.38%) | 0 | 0 | 0 | 0 | 0 | 0 | 0 |

Supplementary Table S1. Results of Ablation experiments.
